# hgtseq: a standard pipeline to study horizontal gene transfer

**DOI:** 10.1101/2022.10.24.513588

**Authors:** Simone Carpanzano, Mariangela Santorsola, nf-core community, Francesco Lescai

**Affiliations:** Department of Biology and Biotechnology, University of Pavia, Pavia, Italy

**Keywords:** horizontal gene transfer, exome, sequencing, microbiome, bioinformatic pipelines

## Abstract

Horizontal gene transfer (HGT) is well described in prokaryotes, it plays a crucial role in evolution, and has functional consequences in insects and plants: less is known about HGT in Humans. Studies have reported bacterial integrations in cancer patients, and microbial sequences have been detected in data from well-known Human sequencing projects. Few of the existing tools to investigate HGT are highly automated. Thanks to the adoption of Nextflow for life sciences workflows, and the standards and best practices curated by communities such as nf-core, fully automated, portable, and scalable pipelines can now be developed. Here we present nf-core/hgtseq, to facilitate the analysis of HGT from sequencing data in different organisms. We showcase its performance by analysing six exome datasets from five mammals. Hgtseq can be run seamlessly in any computing environment and accepts data generated by existing exome and whole-genome sequencing projects: this will enable researchers to expand their analyses into this area. Fundamental questions are still open, about the mechanisms and the extent or the role of horizontal gene transfer: by releasing hgtseq we provide a standardised tool which will enable a systematic investigation of this phenomenon, thus paving the way for a better understanding of HGT.

## 1. Introduction

While whole-genome sequencing (WGS) is becoming the sequencing strategy of choice thanks to the progressively lower prices, whole exome sequencing (WES) remains one of the leading technologies for studying coding regions of large genomes, and for clinical applications in Humans. Most of the projects usually are carried out by aligning the raw reads to the host genome, while unmapped reads are discarded: the discovery by Samuels and colleagues [1] that unmapped reads in Human sequencing data contain a certain number of microbial reads, defined by the authors “a lost treasure”, changed somehow the perspective, and raised the question whether horizontal gene transfer (HGT) occurs in Humans.

Horizontal gene transfer refers to the non-vertical transmission of genetic material between different organisms, allowing for the acquisition of novel traits, including antibiotic resistance, capacity to rapidly adapt to new environments or use of new food sources [2]. HGT is relevant to the variation of both prokaryotic and eukaryotic genomes [3], resulting in some evolutionary trajectories which can be best described by reticulated networks rather than a typical phylogenetic tree [4–7].

HGT is a crucial evolutionary force in archaea and bacteria, enabling such organisms to acquire new genetic functions via transduction, conjugation and direct transformation by exogenous DNA [8,9].

The proportion of genes with signatures of HGT into eukaryotic nuclear genomes is considered usually lower than in prokaryotes [3]. Recently, a work by Li et al. [10] (not peer reviewed yet) suggests putative HGTs is however widespread also in eukaryotes. Different and more complex mechanisms underlie HGT events in eukaryotes due to the presence of a nucleus, the separation of somatic and germline tissues, as well as genome surveillance mechanisms which prevent or minimise the HGT material integration. The evolutionary and functional significance of the prokaryotes-to-eukaryotes HGT events continues to be debated [11] and the molecular mechanisms driving these events are not entirely clarified. However, the transfer of material from chloroplasts and mitochondria to their eukaryotic hosts [12] is known to exist as “nuclear mitochondrial transfers” (NUMTs) or “nuclear plastid transfers” (NUPTs). The proposed molecular events underlying these phenomena [13,14] might offer some understanding of HGTs in higher organisms. Wolbachia, an obligated endosymbiotic bacterium living in the germline of its animal, insect and nematode hosts is another example of HGT between prokaryote endosymbionts and their eukaryotic hosts. The proximity of this endosymbiont to the germ cells of its hosts makes it an ideal donor for heritable HGT to eukaryotic genomes. Several hypotheses have been proposed for the proximity-driven transfer of material, but robust evidence is still missing. Moreover, the constant exposure of multicellular organisms to microbial species (parasites, endosymbionts, or the microbiome) challenges the capacity to distinguish between true HGTs, detectable in genome sequencing data from the host, and contaminants acquired during sample collection in large sequencing projects. Hongseok Tae et al. [15] analysed 150 datasets from 1000 Genomes Project and their results indicated that each sequencing centre had specific contaminating signatures defined as “time stamps”. More importantly, DNA extraction kits have been proved to play an important role in affecting the results of HGT analyses, since some DNA extraction methods appear to be prone to pull down specific DNA sequences from selected microbial genera. This issue has been reported to impact especially the discovery of low abundance microbial species [16]. A confirmation of this finding has been provided by Salter and colleagues [17], who conducted an in-depth study on the presence of contaminants related to ordinary extraction kits, after detecting sequences of microbial species in negative controls during PCR. They provided evidence of a specific bacterial signature associated with each of them. Taking advantage of this information, we curated a list of genera to be flagged as potential contaminants and therefore be handled carefully or filtered out entirely during the analysis.

These large sequencing studies are a key focus of interest, because they provide a window into potential HGT events in Humans. Here, horizontal gene transfer would take place mostly in somatic cells: in this case the newly acquired sequences are not transmitted to the next generation but could induce mutations contributing to cancer or autoimmune diseases [18,19]. Riley and colleagues [20] reported the integration of bacterial sequences into samples of Human acute myeloid leukaemia as well as into Human adenocarcinoma samples from the stomach. Cancer is known to be characterised by genomics instability, which might suggest tumour cells to be more permissive to HGT events: several studies have been carried out to confirm this phenomenon [21]. The indication however that other sequencing studies, carried out in control individuals, display a similar presence of microbial-derived sequences in unmapped reads suggests that HGT events might be more widespread than initially thought.

The measure of rates and patterns of HGT loci in metazoan relies on the methods used to identify HGT candidates in genomes and transcriptomes [22]. Parametric methods, including gmos [23], GeneMates [24], ShadowCaster [25] and nearHGT [26], are aimed at finding genetic loci distinguished from their new host genome either by GC content, oligonucleotide spectrum, DNA structure modelling or based on their genomic context. Explicit and implicit phylogenetic methods compare tree topologies and analyse distances between genomes [27], respectively. Examples of these are ShadowCaster and HGT-Finder [28]. These methods have developed rapidly in the last years fuelled by the availability of large datasets of complete eukaryote genomes from next-generation sequencing (NGS). However, many of these either lack implementation in standard automated workflows or cannot carry out the analyses from the raw data up to all the down-stream HGT steps. In many cases they do not scale with increasingly large datasets or cannot be easily implemented in any environment [29]. To date, few highly automated pipelines for revealing HGT events are available. These include the Daisy pipeline by Trappe et al. [30] and its updated version DaisyGPS [31] for the detection of HGTs directly from NGS reads. They are mapping-based tools and require sequencing data from the HGT host organism and reference sequences of both host and donor genomes. Baheti et al. [32] developed the HGT-ID workflow, which detects the viral integration sites in the Human genome. This workflow is fully automated from the raw reads to the viral integration site detection and downstream visualisation. It is however limited to the analysis of viral HGT. Other existing methods for identifying viral integration sites are VirusSeq software [33], ViralFusionSeq [34], VirusFinder [35], and VirusFinder2 [36] but these are also limited to the investigation of viral integration events. Finally, methods for identification of non-Human sequences in both genomic and transcriptomic libraries of Human tissues include PathSeq [37] and its updated implementation GATK PathSeq [38].

Despite the availability of these solutions, we believe there is still a need for a user-friendly automated pipeline, based on a few fundamental features: accessibility, reproducibility, portability and scalability [39]. Accessibility is a fundamental value, to ensure the widest range of researchers can use the pipeline, in an easy and understandable way. Reproducibility, which is at the basis of the scientific method, requires a version-controlled and documented approach to coding, which ensures the same results are always produced with the same inputs. Portability and scalability are now assuming an ever increasing importance, and can be translated into the capacity to execute the pipeline in any computing environment without major changes to the code, and scale well with the size of the input data.

The introduction of Nextflow [40], an increasingly popular workflow framework, allows the implementation of the above-mentioned values. Nextflow is now largely adopted within research and industry contexts in life science, because of its features and flexibility. This domain-specific language relies heavily on container engines, in particular Docker and Singularity, which ensure all software dependencies are resolved regardless of the environment where the pipeline is executed; this also means the code can be ported seamlessly anywhere. Most importantly, one community of Nextflow users, called nf-core [41], offered the opportunity to converge around specific standards both in terms of coding style and conventions, as well as best practices for the use of bioinformatic tools widely adopted in life sciences and beyond. There has been an investment in the modularity of the pipeline’s composition, which allows code-sharing among those implementing the same tools in different workflows, thanks to the adherence to those best practices. Similarly, Biocontainers are used [42] in most cases. This ensures an even wider standardisation of containers, their design and version-control.

In this context and in line with those values and standards we present here hgtseq, a fully automated pipeline to detect and analyse microbial sequences in unmapped reads from host organisms, and investigate potential evidence of horizontal gene transfer events.

## 2. Results

The hgtseq pipeline has been developed using 21 modules from the shared nf-core repository and 4 “local” modules, as described in materials and methods. 329 commits have been made to the pipeline repository, excluding contributions to shared repositories, in order to compose the pipeline. The release 1.0.0 (doi: 10.5281/zenodo.7244735) accepts either a FASTQ with raw paired-end reads from Illumina sequencing as input, or an already aligned paired-end BAM file (Figure 1).

**Figure 1.**
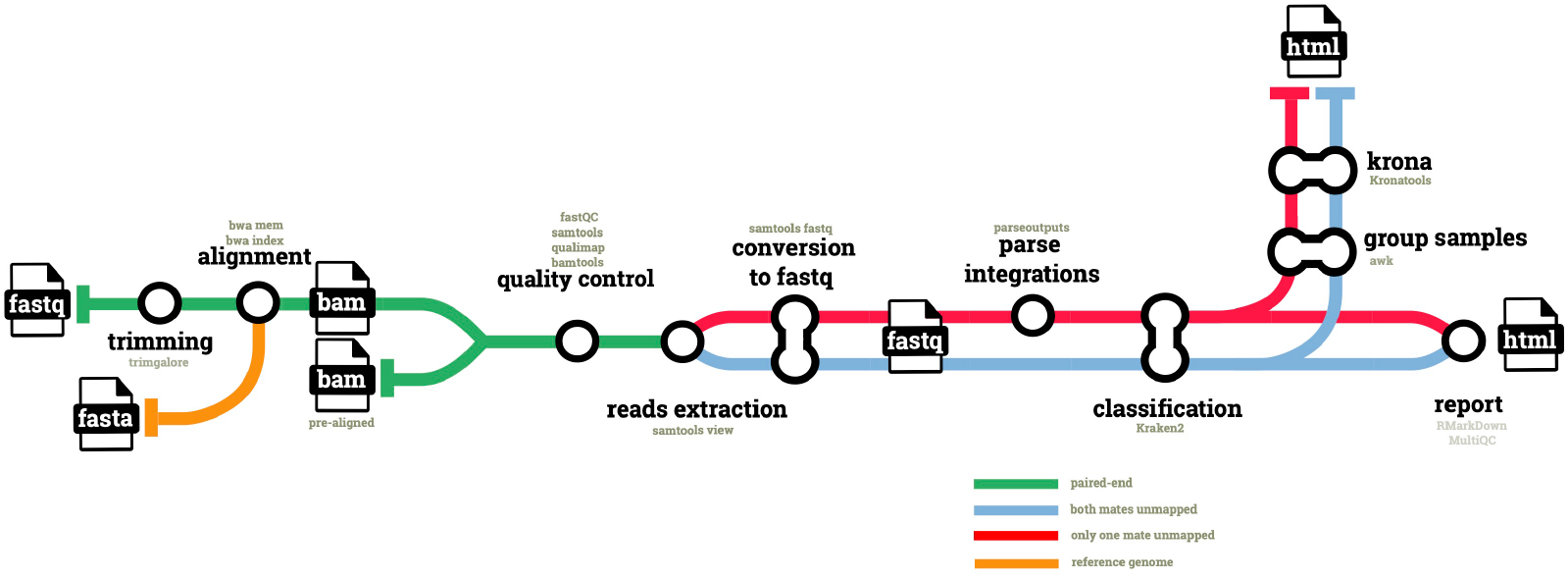
Diagram of the hgtseq pipeline. The figure shows the key steps and input/output files composing the pipeline. Different colours indicate the processing of different data types, such as unmapped reads extracted from the aligned BAM files, which are categorised using samtools flags, depending on whether both mates are unmapped, or just one of the two mates is unmapped.

Raw reads are first trimmed for quality and to remove Illumina adapters: the resulting high quality reads are aligned to the host genome, which is defined by its identifier in the iGenomes repository (https://support.illumina.com/sequencing/sequencing_software/igenome.html) for seamless download, and via NCBI taxonomic identifier. Pre-aligned BAM files are then processed in parallel to extract two categories of reads, via their SAM bitwise flags. With bitwise flag 13, we extract reads classified as paired, which are unmapped and whose mate is also unmapped (i.e. both mates unmapped). With bitwise flag 5 we extract reads classified as paired, which are unmapped but whose mate is mapped (i.e. only one mate unmapped in a pair). In both cases we use flag 256 to exclude non-primary alignments. Both categories are classified using Kraken2 as described in methods. The second category, i.e. unmapped reads whose mate is mapped, provides the opportunity to infer the potential genomic location of an integration event, if confirmed. To this end, the pipeline uses information available for the properly mapped mate in the pair: for this category of reads, the pipeline parses the genomic coordinates of the mate from the BAM file and merges them with the unmapped reads classified by Kraken2.

Finally, host-classified reads are filtered out and the data are used to generate Krona plots and an HTML report with RMarkdown as described in the methods (Figure 1). A full set of QC metrics is available for all steps of the pipeline, and an additional QC report is compiled using MultiQC.

The infrastructure provided by the nf-core community allows the user to access the most important information about the usage and the configuration parameters via a website (https://nf-co.re/hgtseq): these pages are automatically updated from the github repository of the pipeline, and render the markdown documentation together with a JSON schema of the pipeline parameters. This website also offers a “launch” button, prominently displayed at the top of the landing page: this feature allows a user to configure all parameters on the page, and generate a simple JSON file to be passed at command line, thus simplifying the execution at the terminal. Alternatively, the launch feature allows the user to access the Nextflow Tower launchpad (https://cloud.tower.nf) and manage the execution on the cloud through a graphical interface.

We have tested the pipeline on 6 different datasets, downloaded as described in materials and methods: we have verified that the pipeline can be run seamlessly in either an on-premise HPC (slurm scheduler), or on the cloud (Amazon AWS). In our system, the execution of all steps on a Human exome (COPD dataset) completed in just about 3 hours (Figure 2A), generating a total number of 150 jobs (Figure 2B). This corresponds to a total of 154.3 CPU/hours and a total memory usage of 1,136.87 GB. The dataset generated a total I/O of 3,742.28 GB read and 2,389.91 GB written.

**Figure 2.**
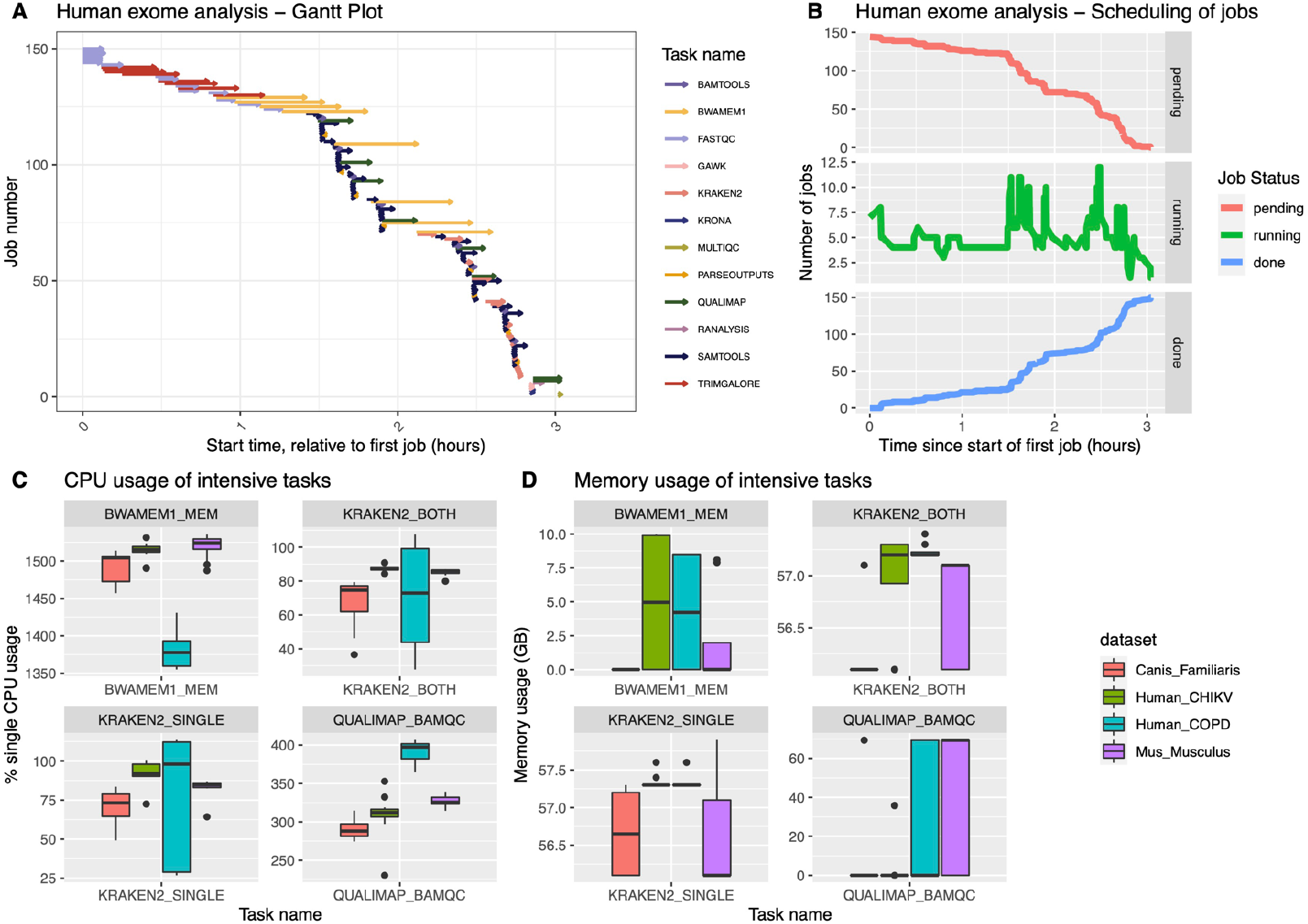
Progress and performance of the pipeline execution. The figure illustrates the execution timeline and the performance of key steps of the pipeline: (**a**) Gantt plot of the analysis of the Human COPD dataset: each arrow indicates the start and the end of the task; the colour code represents the category of the analysis or the main software used in that step; (**b**) The plot represents the progress of the analysis in a local HPC cluster, with each panel representing the current status of each task at the corresponding time point; (**c**) The whiskers plot represent the usage of CPU during the analysis of four of the test datasets, for tools requiring most resources in the pipeline: “single” and “both” in the plot refer respectively to those reads with only one unmapped mate in the pair, and those reads where both mates are unmapped; (**d**) Similarly to plot **c**, the graphical representation shows the memory consumption of the four intensive steps we selected.

The pipeline includes a few well-known compute intensive tasks, such as BWA-MEM for the alignment, Kraken2 for taxonomic classification and Qualimap for the QC of the BAM files, as described in the methods section. BWA showed the ability to handle well multi-threading in different datasets (four of them showed in Figure 2C), while Kraken2 mostly uses 1 core during the classification process. In terms of memory usage, as expected, both Kraken2 and Qualimap use the most amount of memory among all processes in this pipeline, reaching 60GB of memory (Figure 2D), particularly in datasets from Human and Dog. The memory usage of Qualimap shows to be related to the size of the BAM files. Kraken2 memory usage instead shows dependency on the chosen database for the analysis.

The execution on AWS, using the AWS-batch API which Nextflow supports, allowed to confirm the portability and reproducibility of the code.

The samples included in the six datasets we analysed for testing, ranged between 49,516,660 in the Bos taurus cohort to 198,822,100 in the Canis lupus familiaris dataset (Table S1). As described above, we analysed two category of unmapped reads separately (i.e. depending whether both mates or only one mate of the pair was unmapped): Human datasets showed a proportion of unmapped reads classified to microbial genera, when compared to the total number of reads in the sample, ranging between 1.04×10^−5^ (Human CHIKV cohort, both mates unmapped) to 2.44×10^−6^ (Human COPD dataset, both mates unmapped - Figure 3 and Table S1). Similar ranges where observed in the other species: the proportion in Bos Taurus ranged between 8.5×10^−5^ (both mates unmapped) to 8.67×10^−5^ (only one mate unmapped); in Canis lupus familiaris was 3.42×10^−8^ for both mates unmapped reads and 7.32×10^−7^ for unmapped reads having their mate mapped; The proportion in Macaca fascicularis was 1.16×10^−5^ for both mates unmapped and 2.14×10^−5^ for one mate only unmapped reads, while in Mus musculus we observed a proportion of 4.91×10^− 6^ for both mates unmapped and 6.21×10^−6^ for one mate only unmapped.

**Figure 3.**
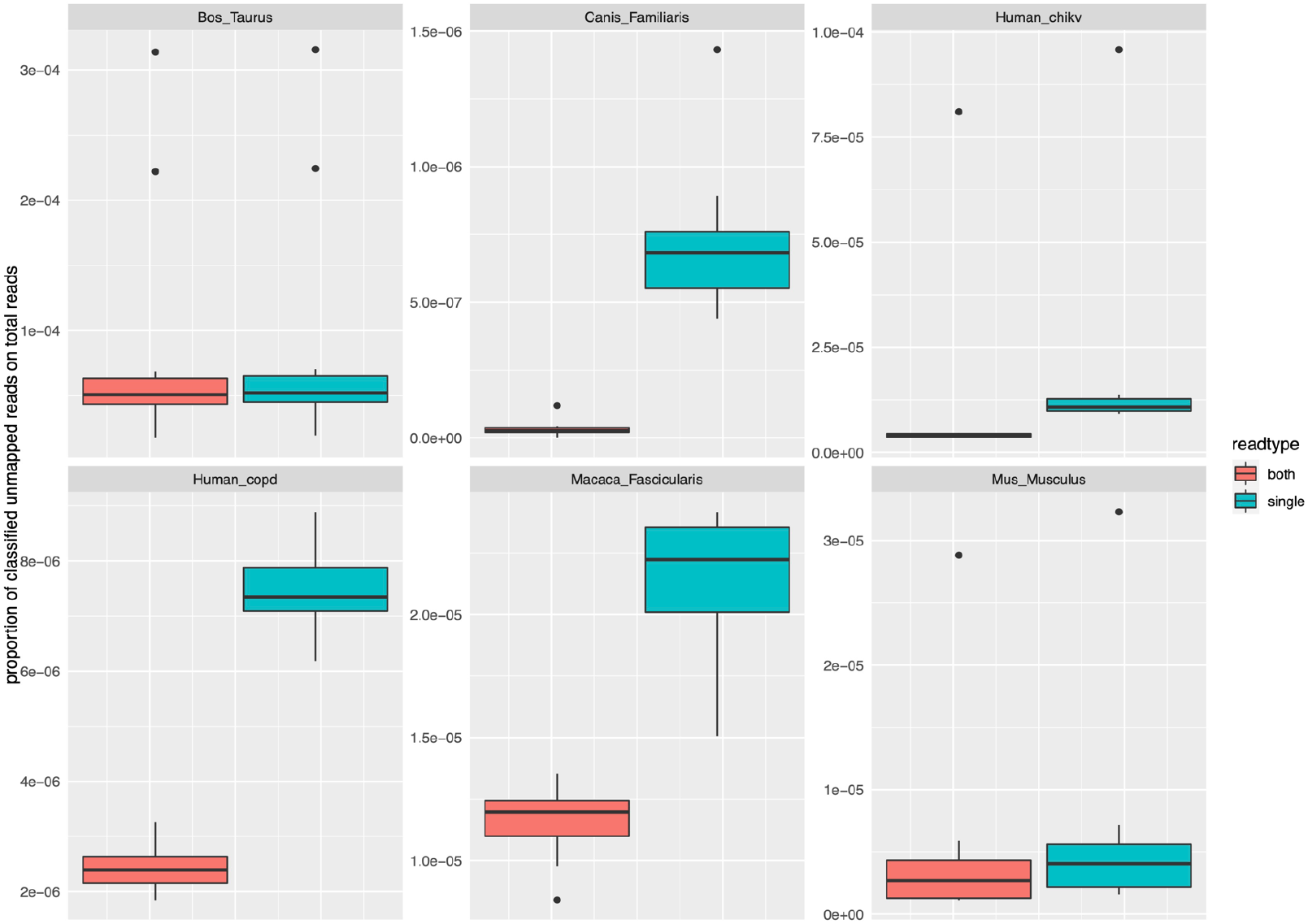
Proportion of reads assigned to microbial genera in different datasets, compared to the total number of reads in the sample. The figure illustrates the proportion of unmapped reads assigned to microbial genera in each of the test datasets. The reads are categorised by type, with “both” indicating those where both mates in the pair are unmapped and “single” those where only one mate in the pair is unmapped.

We then set out to understand the proportion of potential contaminants in these results, by assigning a flag to the classified reads, depending on whether the genus of their assignment belonged to the list of genera we compiled, as described in materials and methods. We assigned the category “flagged” if the genus was included in the list, but belonged to genera which might include species known to be associated with Humans. We assigned the category “blacklisted” if the genus belonged to any other genera in the list. We also stratified this analysis using the classification score we calculated in the final step of the pipeline analysis. This analysis (Figure S1) shows that in most of the datasets the proportion of classified reads belonging to potentially contaminant genera is higher among those reads with a lower classification score (Mus musculus, Macaca fascicularis, Human in the CHIKV dataset, Canis lupus familiaris). This cannot be however observed in all cases, as the Human COPD dataset shows a higher proportion of flagged genera in reads with higher classification scores, and the results in the Bos taurus dataset show an unexpectedly high proportion of flagged reads overall.

In order to explore these results, we then analysed in particular the classified reads belonging to the category single-mapped (i.e. those unmapped reads whose mate in the pair is mapped). The vast majority of the reads have been assigned to bacterial organisms, and we therefore compared the krona plots generated on those reads from each of the different datasets we processed, after filtering out blacklisted genera (Figure S2). A qualitative inspection of the plots shows clear differences between the datasets of Bos taurus and Mus musculus, while a number of similarities can be observed in primates (Human COPD and CHIKV, and Macaca fascicularis).

## 3. Discussion

Since the first identification of a significant presence of non-Human sequences in Human data [1], and a large-scale follow-up in different sequencing data [15], this quite intriguing topic has been poorly investigated. Horizontal gene transfer events between bacteria and eukaryotes have been described, and several molecular mechanisms controlling these events have been proposed [2]: while those phenomena in plants and invertebrates are documented in detail, much less in known about horizontal gene transfer events in mammals, which might explain the initial findings of Samuels and colleagues. The available data support the existence of horizontal gene transfer events in mammals, including Humans, and call for a renewed effort in investigating this phenomenon in a systematic way, exploiting the wealth of sequencing data already available in a variety of species and conditions. The main obstacle to this, is therefore represented by the availability of tools to enable these analyses at scale, with any type of data and in the widest variety of computing environments available to researchers. Here we documented our work to release a standard pipeline to investigate the presence of microbial DNA in mammals, characterised by ease of use, portability and scalability.

The first critical choice we made, in order to achieve this goal, has been to develop the code by using Nextflow. The nature of this domain specific language, agnostic to the code or the software being used, makes it both flexible and capable to accommodate a wide range of choices, in terms of tools for the pipeline. Nextflow supports most schedulers for on-premise high performance computing clusters, and all cloud infrastructure vendors: a pipeline developed with this workflow engine can therefore be run virtually anywhere, with just little changes to the configuration files. The second fundamental choice we made, was to develop the pipeline within the framework of nf-core: besides the standards and best practices, the community also developed an infrastructure which allows the code to be continuously tested, reviewed by a large number of people, and shared among different pipelines. Common choices are usually discussed among scientists from different disciplines and institutions. Additionally, a strong focus on documentation and the availability of a website to configure pipeline parameters and launch the pipelines on the cloud, makes the nf-core infrastructure a best-in-class choice for life science applications to be used at scale. Hgtseq benefits from this crucial environment and facilities, and has therefore the potential to push this area of research forward. The pipeline accepts both FASTQ and BAM/CRAM files as input: this has been a strategic choice, meant to allow executing hgtseq downstream to a wide range of routine pipelines, particularly in Human studies. This way, we meant to target research groups who primarily study Human or other host genomes, but do not look for non-host reads in their data. With this in mind, we hope to offer additional opportunities to tap into a wealth of data available, which can be used to answer the fundamental questions still open. Additionally, the modularity the pipeline has been designed with provides further opportunities to include selected features from solutions implemented in other valuable tools, thus expanding the capabilities in further releases.

In order to demonstrate the functionality of this pipeline, we chose a wide array of species to run the analyses on. Besides two different Human datasets, we decided to use other mammals whose genome is well curated, such as Bos taurus, Mus musculus and Canis lupus familiaris. We also chose Macaca fascicularis, to provide an additional insight of the pipeline performance on sequencing data from another primate. For all these species, we chose to analyse exome data: this was meant to mirror the initial analyses by Samuels and colleagues [1], but also to test the capability of hgtseq to detect this phenomenon within the coding regions of other mammalian species.

In terms of performance, we have been able to demonstrate that the pipeline can handle a wide variety of sequencing data, and process datasets with more than 100 million reads per sample in about 3 hours. The speed of the analysis will naturally depend on the computing environment where it is run, and we provide here an indication on the resources needed in terms of CPUs and memory by the most resource-consuming tools in this pipeline, such as BWA, Kraken2 and Qualimap. Users will be able to plan their analyses based on this information. Changes to the configuration can be easily introduced by creating new config files for the infrastructure to be used at the time of analysis.

The analysis we present here, has the limited scope of testing the computational performance of the pipeline we introduce, and showing its basic functionality. Within the limited size of the datasets we used, we have been able to confirm that the presence of microbial sequences in exome data is not limited to Human exomes, but exists to a similar extent also in other mammalian species. In particular, in this work we are able to confirm the findings of Tae et al. [15] in terms of abundance of microbial species in this kind of data. When evaluating these findings though, it is quite crucial to assess the extent of a bias in measuring the abundance of unmapped reads classified to microbial taxonomies. A source of bias might be the potential contamination of the sequencing data. Within the pipeline we have provided two ways to control for this: first, we have curated a list of genera which are reported over the years to be known contaminants of DNA extraction kits. This list, accompanied by a review of the literature used as a source, can be embedded into the analyses and be used to filter the results. When analysing Human data, we have further categorised this list, by highlighting those microbial genera which might include species known to be associated with a Human host: the user can therefore decide whether those results should be filtered out, or marked for a closer inspection. Additionally, we have coded an evaluation score into the results report, to further stratify the strength of Kraken2 taxonomic assignment of each read: this score provides a second tool to filter the results and fine tune the stringency of the analysis. In our tests, we have shown that using this score we can control the abundance of genera flagged for contamination. We have also highlighted the extent of variability in sequencing quality among different datasets.

In conclusion, we believe there is an urgent need to systematically investigate the extent of horizontal gene transfer phenomena in mammals. There are still several open questions to be answered, about the molecular mechanisms controlling these events, or the extent mammal genomes are affected by them, and what the source of microbial material might be. While a much larger analysis is needed to address these questions, we are offering the scientific community a standardised, portable and scalable tool to carry out a systematic investigation of these events in a wide range of hosts, thus paving the way for new potential breakthroughs in this area of research.

## 4. Materials and Methods

### 4.1 Datasets

Six exome datasets from different species have been chosen to test the pipeline on a variety of conditions. Two datasets from Human exomes have been downloaded: 9 samples from a Chinese population affected by chronic obstructive pulmonary disease (COPD), available from SRA (BioProject accession: PRJNA785331); 12 samples from Brazilian individuals with Chikungunya virus (CHIKV) infection, available from SRA (BioProject accession: PRJNA680041). Additionally, we have used 10 samples from a population of whole exome sequence from 69 male founder mice (BioProject accession: PRJNA603006), 12 samples from a dataset of Macaca fascicularis exomes (BioProject accession: PRJNA685531), 10 samples from an exome dataset of Bos Taurus (BioProject accession: PRJNA479823) and 10 samples of Poodle breed from a project of exome sequencing in three dog breeds (Canis lupus familiaris, BioProject accession: PRJNA385156).

### 4.2 Alignment and read extraction

When FASTQ files are used as input, the pipeline first uses TrimGalore (https://www.bioinformatics.babraham.ac.uk/projects/trim_galore/) to trim the reads by quality and to remove any adapters (default options, plus ‘--illumina’). Trim Galore combines both Cutadapt (https://github.com/marcelm/cutadapt/) and Fastqc (https://www.bioinformatics.babraham.ac.uk/projects/fastqc/) to perform both types of trimming in the same step. These processed reads are then mapped to the host reference genome using BWA [43], with default parameters. These steps are automatically skipped if the user provides pre-aligned BAM files as input. According to standard practice, the aligned BAM files are then sorted by chromosome coordinates and indexed.

Using ‘*samtools view*’ [44], we extract two different types of reads pairs identified by their SAM bitwise flag: each read type is then processed separately, according to their different properties. Specifically, to extract the first category of data we apply the command ‘*samtools view -b -f 13 -F 256*’ to select those reads with bitwise flag 13 and therefore extract those in a pair where both mates do not map to the host reference (flag properties: read paired, read unmapped, mate unmapped); we define them as *both_unmapped*. To extract the second category of data, we apply the command ‘*samtools view -b -f 5 -F 256*’ to select reads with a bitwise flag 5, i.e. to extract those unmapped reads in a pair whose corresponding mate is mapped (flag properties: read paired, read unmapped); we define them as *single_unmapped*. As can be seen in these commands, by applying the option ‘-F’ in both cases we use the bitwise flag 256 to exclude any non-primary alignments.

A local module named *parseoutputs*, which contains two combined commands ‘samtools view | cut -f 1,3,8’ allows the extraction from the *sigle_unmapped* reads of mapping information of their mapped mate: sample name, chromosome and exact position of the mate. These data are merged with the taxonomic classification of the unmapped reads in the RMarkDown report, for further analyses of potential integration sites.

### 4.3 Reads classification

In order to perform the taxonomic classification of the unmapped reads extracted as described above, the pipeline uses Kraken2 [45], a k-mers based algorithm that provides a high-accuracy classification by matching each k-mer to the lowest common ancestor. Reads are first converted into a FASTQ format to accommodate Kraken2 input requirements.

We built a custom database with all available libraries in Kraken2 (archea, bacteria, plasmid, viral, fungi, plant, protozoa, human) by using the command ‘*kraken2-build – download-library*’. Additionally, we downloaded from ENSEMBL the genome of other host species to be used in this analysis such as Canis familiaris, Bos taurus, Macaca fascicularis and Mus musculus. We use an in-house python script to reformat the FASTA header of the genome sequences, according to Kraken2 specification. Finally, we added the sequences to Kraken2 library with the command ‘*kraken2-build – add-to-library*’, and build the database. We then run Kraken2 on the two categories of unmapped reads with default settings, and collect the default output (report, and reads assignment).

### 4.4 Reporting

The QC report is generated by MultiQC [46], by aggregating data from multiple sources within the pipeline, and in particular FastQC (https://www.bioinformatics.babraham.ac.uk/projects/fastqc/), statistics generated by samtools [44], bamtools stats (https://github.com/pezmaster31/bamtools) and Qualimap [47].

The analysis report is generated via a specific process in the pipeline, which aggregates results from all samples and executes a parameterised RMarkdown [48] to create tables and plots.

In order to provide a measure of the evidence each read classification is based upon, we have added a classification score to the analysis that generates the report on the datasets. The R code parses the k-mer information from Kraken2 output, and counts: (a) the total number of k-mers analysed in each read (*tot_kmers*), (b) the sum of all k-mers supporting the classified taxonomic ID, which is chosen by Kraken2 as the one with the highest number of supporting k-mers (*max_kmers*), (c) the total number of classified k-mers in the read, i.e. with taxonomic ID not zero (*sum_class_kmers*). A classification score (*taxid_score*) is then provided as the ratio *max_kmers/tot_kmers*; an additional score is also provided as the ratio *max_kmers/sum_class_kmers*, i.e. indicating the proportion of the assigned taxonomic ID over other classified taxonomic IDs.

Besides this RMarkDown report the pipeline generates two multi-layered interactive pie charts made with Kronatools [49]. For convenience, we wrote a local module that groups together all samples to make only one chart from each of the two read categories. Using the command ‘ktImportTaxonomy -t 3’ this tool parses only the taxa from each read.

### 4.5 Pipeline composition

In order to compose the pipeline, we have used version 2 of the domain-specific language (DSL2) Nextflow [40]. This version of the workflow language allows the combination of multiple independent scripts (defined as modules), each defining functions, processes and/or workflows. The nf-core community has released standard guidelines [41], identifying modules as atomic scripts, where one process uses one single tool or tool function. This approach makes it possible to share nextflow modules within the community across multiple users and pipelines. In the process of designing the pipeline, the modules of general use have been therefore contributed to a shared nf-core repository first (krona, kraken2, samtools utilities, bamtools stats), and they have subsequently been included in the hgtseq repository. Those scripts specifically designed to carry out tasks within hgtseq, have been committed to the hgtseq repository as “local” modules. Each module uses standardised containers, curated by the community Biocontainers (https://biocontainers.pro).

We have used the pipeline template developed by nf-core and the nf-core tools (https://nf-co.re/tools/), released to facilitate pipeline composition according to community standards. These include continuous integration (CI) tests run using the tool *pytest* which are triggered by github actions at commits push or pull requests. An extensive array of CI tests is being run, using three containers/package managers (docker, singularity, conda) on the 2 key use cases of the hgtseq pipeline, i.e. with FASTQ input, and with BAM input.

Additionally, the design includes a full-size test run at pipeline release on AWS and using the Human COPD samples (BioProject accession: PRJNA785331) as input.

## Supplementary Materials

The following supporting information can be downloaded at: www.mdpi.com/xxx/s1, Figure S1: Relative abundance of genera indicated as potential contaminants; Figure S2: Krona plots of single-unmapped reads in all datasets; Table S1: Percentage of reads assigned to microbial species in different datasets.

## Author Contributions

Conceptualisation and methodology: FL and SC; Software and validation: SC, FL and the nf-core community wrote and reviewed the code; Formal analysis: FL and SC; Writing, original draft, reviewing and editing: SC, FL and MS; Resources: the nf-core community provided the infrastructure to facilitate access to documentation and graphical user interfaces.

## Data Availability Statement

The code is available at https://github.com/nf-core/hgtseq. The release 1.0.0 of the pipeline can also be referenced with doi://10.5281/ zenodo.7244735.

