## Supplementary Materials for "hgtseq: a standard pipeline to study horizontal gene transfer"

**Figure S1**

Relative abundance of genera indicated as potential contaminants

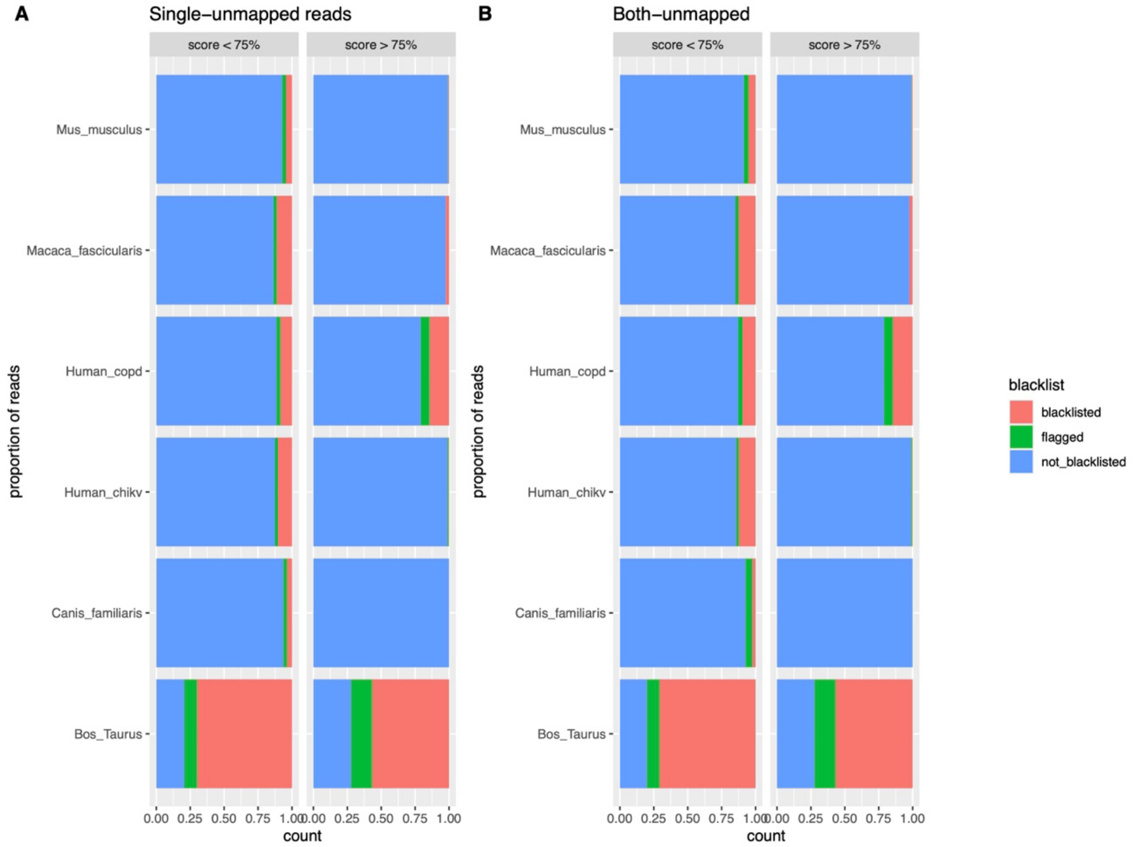


The figure illustrates the proportion of reads classified to microbial organisms, and assigned to genera listed among potential contaminants. The plot represents all test datasets and is stratified by Kraken2 classification score (either above or below 75%) and by read type, i.e. “single-unmapped” when referring to reads whose mate is mapped, and “both-unmapped” when referring to reads where both mates in the pair are unmapped. Two flags are reported among potential contaminants: “not_blacklisted” (in blue) indicates those taxonomic assignments not belonging to genera considered contaminants, “blacklisted” (red) refers to genera flagged as potential contaminants, while “flagged” (green) refers to genera listed as potential contaminants but including species known to be of potential interest for Human infections.

**<u>Figure S2</u>**

Krona plots of single-unmapped reads in all datasets

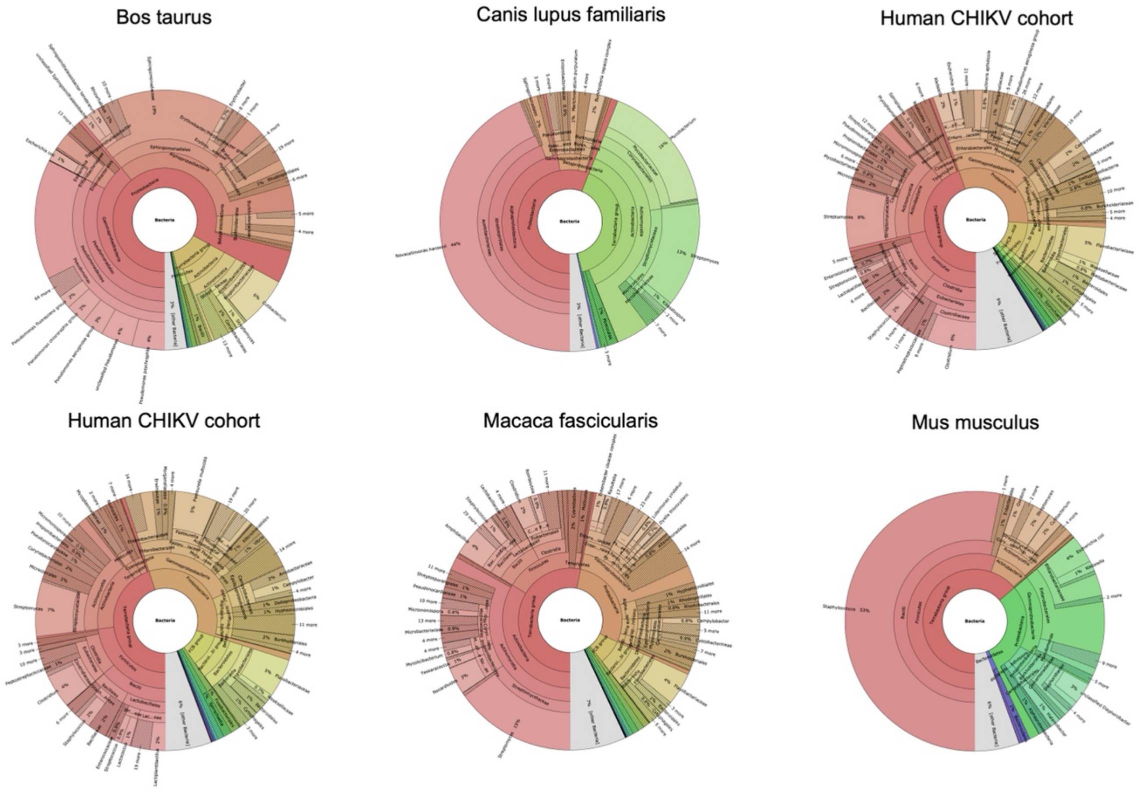


The figure combines krona plots generated in all datasets from unmapped reads whose mate in the pair is mapped (“single-unmapped”), after filtering out blacklisted genera, as represented in Supplementary Figure 1.

**<u>Table S1</u>**

Percentage of reads assigned to microbial species in different datasets

| Dataset | Average reads per sample | Unmapped and microbial classified read type | Average proportion of microbial classified unmapped reads over total sample reads |
|---|---|---|---|
| Bos_Taurus | 49,516,660 | both mates unmapped | 8.50E-05 |
| | | only one mate unmapped | 8.67E-05 |
| Canis_Familiaris | 198,822,100 | both mates unmapped | 3.42E-08 |
| | | only one mate unmapped | 7.32E-07 |
| Human_chikv | 160,862,500 | both mates unmapped | 1.04E-05 |
| | | only one mate unmapped | 1.81E-05 |
| Human_copd | 117,495,800 | both mates unmapped | 2.44E-06 |
| | | only one mate unmapped | 7.45E-06 |
| Macaca_Fascicularis | 76,028,120 | both mates unmapped | 1.16E-05 |

| | | | |
|---|---|---|---|
| | | only one mate unmapped | 2.14E-05 |
| Mus_Musculus | 100,482,900 | both mates unmapped | 4.91E-06 |
| | | only one mate unmapped | 6.21E-06 |

The table represents the data underlying Figure 3 in the main text, and reports the average number of reads per sample in each of the test dataset, as well as the proportion of unmapped reads assigned to microbial organisms, by read type.
